# Alanine scanning mutagenesis of the MEDI4839 (Suvratoxumab) epitope reduces alpha toxin lytic activity *in vitro* and *S. aureus* fitness in infection models

**DOI:** 10.1101/323642

**Authors:** C Tkaczyk, E Semenova, Y.Y Shi, K Rosenthal, V Oganesyan, P Warrener, C.K Stover, B.R Sellman

## Abstract

Alpha toxin (AT) is a cytolytic pore-forming toxin that plays a key role in *Staphylococcus aureus* pathogenesis, consequently extensive research was undertaken to understand the AT mechanism of action and its utility as a target for novel prophylaxis and treatment strategies against *S. aureus* infections. MEDI4893 (Suvratoxumab) is a human anti-AT IgG1 monoclonal antibody (mAb), which targets AT and is currently in Phase 2 clinical development. As shown previously, the MEDI4893-binding epitope on AT is comprised of the highly conserved amino acid regions 177-200 and 261-271, suggesting these amino acids are important for AT function. To test this hypothesis, and gain insight into the effect mutations in the epitope on AT neutralization by MEDI4893, nine MEDI4893 contact residues in AT were individually mutated to alanine. Consistent with our hypothesis, 8 out of 9 mutants exhibited >2-fold loss in lytic activity resulting from a defect in cell binding and pore formation. MEDI4893 binding affinity was reduced >2-fold (2 – 27-fold) for 7 out of 9 mutants and no binding detected for W187A mutant. MEDI4893 effectively neutralized all of the lytic mutants *in vitro* and *in vivo*. When the defective mutants were introduced into a *S. aureus* clinical isolate, the mutant-expressing strains exhibited less severe disease in mouse models and were effectively neutralized by MEDI4893. These results indicate the MEDI4893 epitope is highly conserved due, in part to its role in AT pore-formation and bacterial fitness, thus decreasing the likelihood for the emergence of mAb-resistant variants.

## Importance

Increasing incidence of antibiotic resistant bacteria and desire to protect the healthy microbiome raises the need for testing alternative strategies of antibacterial therapy such as immunotherapy. We previously generated the human monoclonal antibody, MEDI4893 (Suvratoxumab), against secreted alpha toxin (AT), a key virulence factor for *Staphylococcus aureus* pathogenesis, currently in clinical testing for prevention of *S. aureus* pneumonia. The AT sequence is well conserved among clinical isolates including the nine MEDI4893 contact residues on AT. To better understand their role on AT function and mAb neutralizing activity, we tested the effect of mutations in the nine contact residues *in vitro* and *in vivo* lytic activity and disease severity. Several mutants exhibited reduced activity *in vitro* and *in vivo*, but none escaped MEDI4893 neutralization. These data demonstrate the conserved MEDI4893 epitope is essential for AT lytic activity and bacterial fitness, therefore providing a hurdle to the emergence of MEDI4893 resistant variants.

## Introduction

The spread of antibiotic resistance along with a better understanding of the adverse effects broad-spectrum antibacterial therapy has on the beneficial microbiome have led to the exploration of alternative approaches to antibacterial therapy including pathogen-specific monoclonal antibodies (mAbs) to prevent or treat serious bacterial infections (1, 2). MEDI4893 (Suvratoxumab) is a high affinity anti-*S. aureus* alpha toxin (AT) mAb currently in Phase 2 clinical development for the prevention of *S. aureus* pneumonia in mechanically ventilated patients colonized with *S. aureus* in the lower respiratory tract (3). Previous studies demonstrated that AT acts as a key virulence factor in numerous preclinical disease models including dermonecrosis, lethal bacteremia and pneumonia (4–7). There is also evidence that AT is important in human disease as high AT expression levels by colonizing isolates was linked to progression to pneumonia in ventilated patients (8) and low serum anti-AT IgG levels correlate with increased risk for recurrent skin infections in children (9). AT exerts its toxic effects by forming pores in target cell membranes, leading to cell lysis at higher toxin levels (10). It also has effects at sub-lytic levels, resulting in disruption of epithelial and endothelial tight-cell junctions, a damaging hyper inflammatory response in the lung and evasion of killing by host innate immune cells (11–13). Alpha toxin is secreted as a soluble monomer that binds a disintegrin and metalloprotease 10, ADAM10, on cell membranes, oligomerizes into a heptameric ring and undergoes a conformational change resulting in transmembrane pore formation in host cells such as monocytes, lymphocytes, platelets, and endothelial and epithelial cells (10, 14).

Active and passive immunization strategies targeting AT have been reported to reduce disease severity in skin and soft tissue infections, lethal bacteremia and pneumonia (4, 5, 15–19). Specifically, MEDI4893*, a non YTE version of MEDI4893, has been shown to reduce disease severity in multiple animal models (13, 17, 20) and to exhibit synergy when administered in adjunctive therapy with standard of care antibiotics (15, 21, 22). MEDI4893 binds with high affinity to a discontinuous epitope on AT (amino acids 177-200 and 261-27) and inhibits pore-formation by blocking toxin binding to target cell membranes (20, 23). Recent studies of diverse *S. aureus* clinical isolate collections (~1250 total) demonstrated that the AT gene, *hla*, is encoded by a majority (>99.5%) of isolates and 58 different AT sequence variants were identified (24–26). In these collections, the MEDI4893 epitope was highly conserved with only 19 isolates having mutations in the epitope. We hypothesized that such a high degree of conservation results from the amino acids (AA) in the MEDI4893 epitope playing a key function in AT lytic activity.

To better understand the role of these AA in the cytolytic mechanism of AT and gain insight into the effect of mutations in AT MEDI4893-binding epitope, the residues on AT that contact MEDI4893 were mutated to alanine. The results from this study indicate the AA in the MEDI4893 epitope are important for toxin function and bacterial virulence and that the mAb is capable of neutralizing toxin molecules with mutations in its binding epitope. Taken together, our observations imply a low potential for emergence of AT variants resistant to neutralization by MEDI4893.

## Results

### Cytolytic activity of AT alanine mutants

The crystal structure of the MEDI4893 antigen-binding fragment (Fab) bound to AT revealed a discontinuous epitope spanning amino acids (AA) 177-200 and 261-271 with direct molecular contacts on the toxin at residues D183, S186, W187, N188, P189, V190, R200, T263 and K266 (**Fig. 1A** **and B**) (23). These residues were highly conserved among ~1,250 diverse *S. aureus* clinical isolates (24–26). Alanine scanning mutagenesis of these 9 contact residues was conducted to determine their role in AT function and to gain insight into the effect these mutations have on MEDI4893 neutralizing activity. Cytolytic activity of AT alanine mutants was first examined on rabbit red blood cells (**Fig. 2A**) and the A549 human lung epithelial cell line (**Fig. 2B**). As shown in Figure 2B, W187A, N188A and R200A mutants exhibited little or no cytolytic activity on A549 cells. All of the mutants, with the exception P189A on A549 cells and D183A on red blood cells exhibited a >2-fold loss in cytolytic activity on both cell types (Fig. 2). When MEDI4893 was incubated with either the WT or mutant toxins (2:1 molar ratio mAb:AT) prior to the assays, the mAb exhibited neutralizing activity (≥95%) against all mutants in the hemolytic assay with the exception of K266A and W187A against which the mAb neutralized 60% and 0% of their activity, respectively (**Fig. 3A**). MEDI4893 neutralized ≥75% of the cytolytic activity of D183A, S186A, P189A, V190A, T263A and K266A on A549 cells (**Fig. 3B**). W187A, N188A and R200A were not tested in the A549 lysis assay because of their greatly diminished lytic activity. These results indicate that AA in the MEDI4893 epitope are essential for AT function and that the mAb neutralizes epitope variants *in vitro*.

**Fig. 1:**
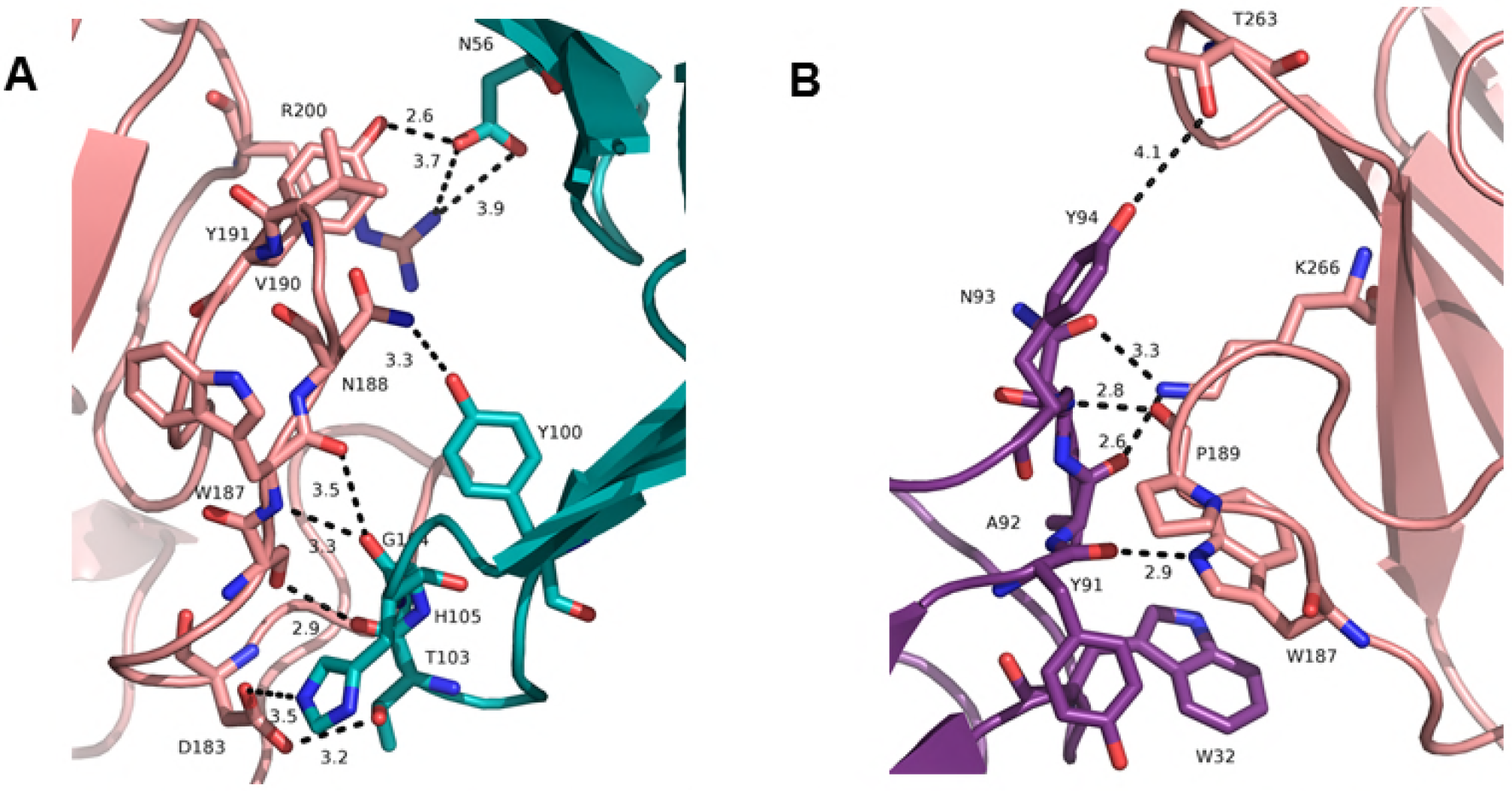
Interface between MEDI4893 Fab HC (green) and AT (pink) (A) and MEDI4893 Fab LC (purple) and AT (pink) (B). Both chain of the Fab interact with AT and create hydrogen bonds (dotted lines). Residue W187 interact with HC through hydrogen bound, and with W32 on LC by π-π stacking interaction.

**Fig. 2:**
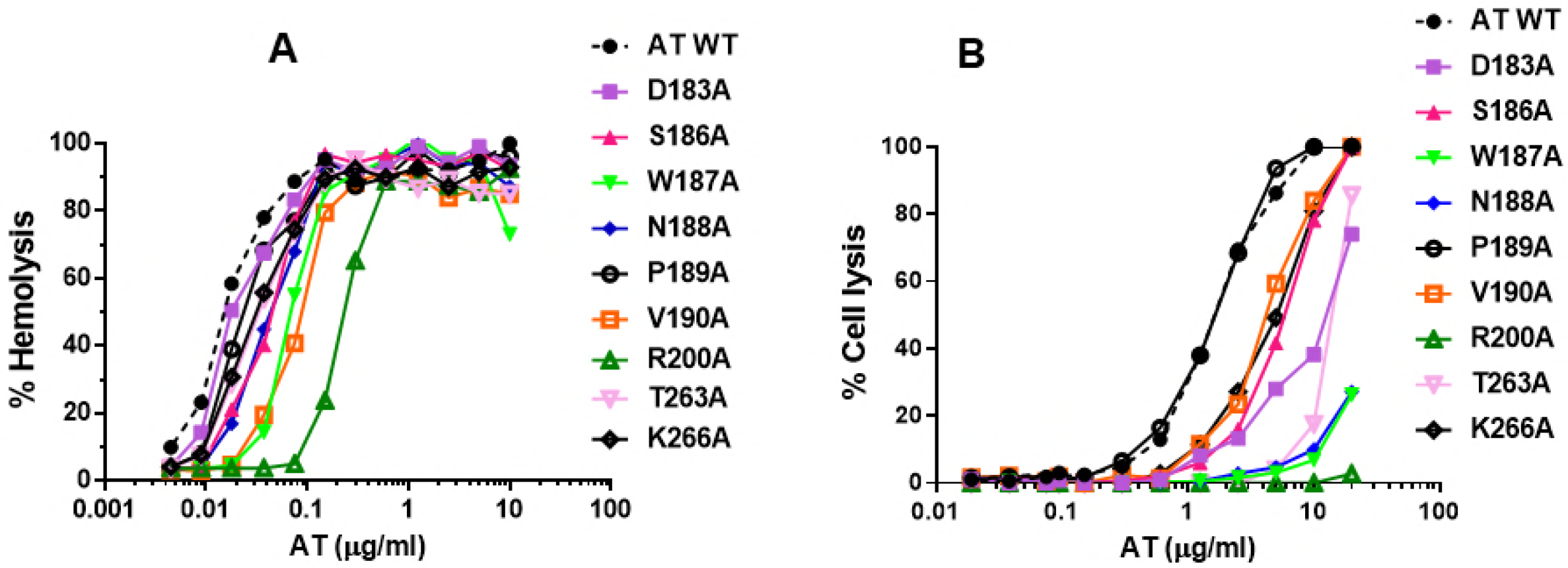
AT mutant lytic activity on rabbit RBC (A) and human A549 cell line (B). **(A)** Washed RBCs were incubated with serial dilutions of nAT or AT mutants (10 to 0.005μg/ml). Hemolysis was measured by the amount of hemoglobin released in the supernatant, and calculated as followed: 100*[(OD_AT_)/(OD_SDS_)]. (B) A549 were incubated with serial dilutions of nAT or AT mutants (20 to 0.01 μg/ml). LDH was measured in supernatants after two hour incubation at 37°C. % cytolysis was calculated as follows: 100*[100-(OD_AT_) / (OD_SDS_)].

**Fig. 3:**
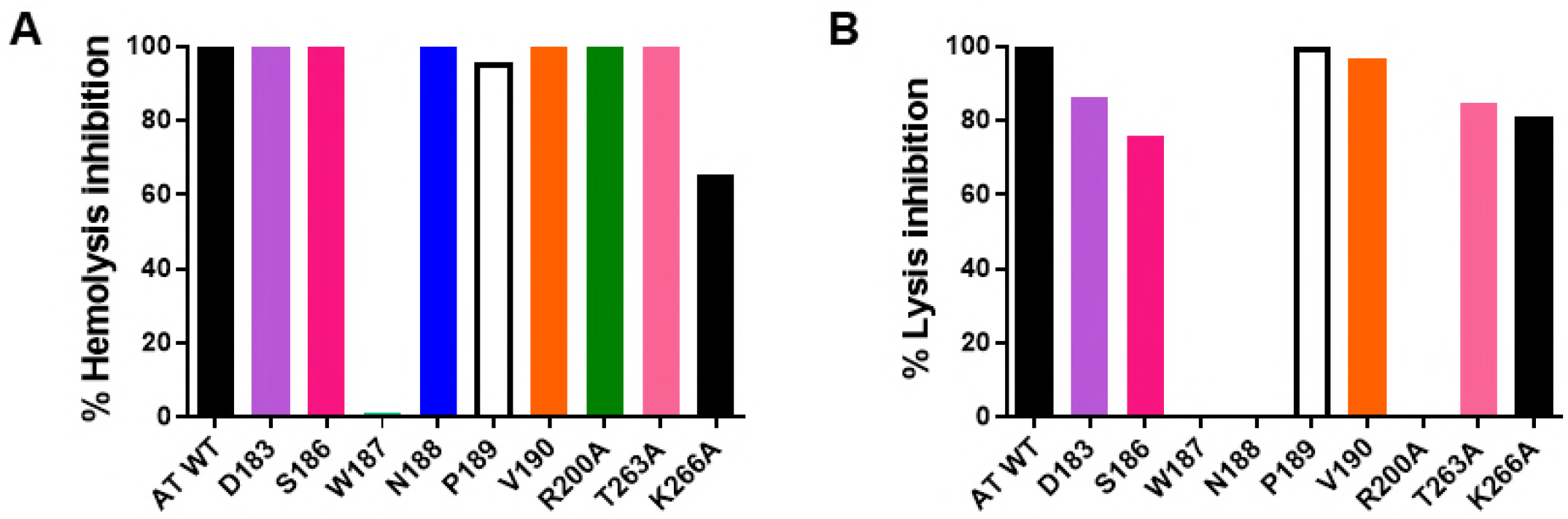
MEDI4893* neutralization ill vitro oil rabbit RBC hemolysis (A) aud A549 lysis (B) AT or mutants were mixed with MED14893* serial dilutions and RBC (0.1μg/ml) (A) or with (B) A549 (20μg/ml). % hemolysis or cell lysis inhibition was calculated as follows: 100*[100-(OD_AT+mAb_) / (OD_AT alone_)]. Data are representative of three independent experiments for AT:MEDI4893* ratio 1:2.

### MEDI4893* reduces AT mutant-induced dermonecrosis

AT is a key virulence factor in *S. aureus* skin and soft tissue infections (5, 6, 27, 28) and intradermal (ID) injection of purified toxin results in dermonecrotic lesions in mice (29). To determine if the *in vitro* lytic activity translated into a similar pattern of activity *in vivo*, the capacity of each alanine mutant to induce dermonecrosis in mice was assessed. WT or mutant toxins were injected ID into Balb/c mice (n=10) and dermonecrotic lesion sizes recorded 24h post-injection. Injections of either P189A or K266A mutants resulted in lesion sizes similar to WT toxin 24h post-injection, whereas D183A, S186A and V190A formed lesions smaller than WT AT. W187A, N188A, R200A and T263A formed little or no detectable lesions (**Fig. 4A** **and Fig. S1**). Results were similar on day 7 except for delayed dermonecrotic lesion formation by D183A, N188A, and T263A (**Fig. S1**). Consistent with its *in vitro* neutralization activity, passive administration of MEDI4893* (10mpk) 24h prior to toxin injection resulted in complete inhibition of lesion formation induced by WT AT and all dermonecrotic AT mutants (**Fig. 4B**). These results indicate the *in vitro* lysis activity remains consistent *in vivo* and provides evidence that MEDI4893 effectively neutralizes *in vivo* AT mutants with AA changes in its binding epitope

**Fig. 4:**
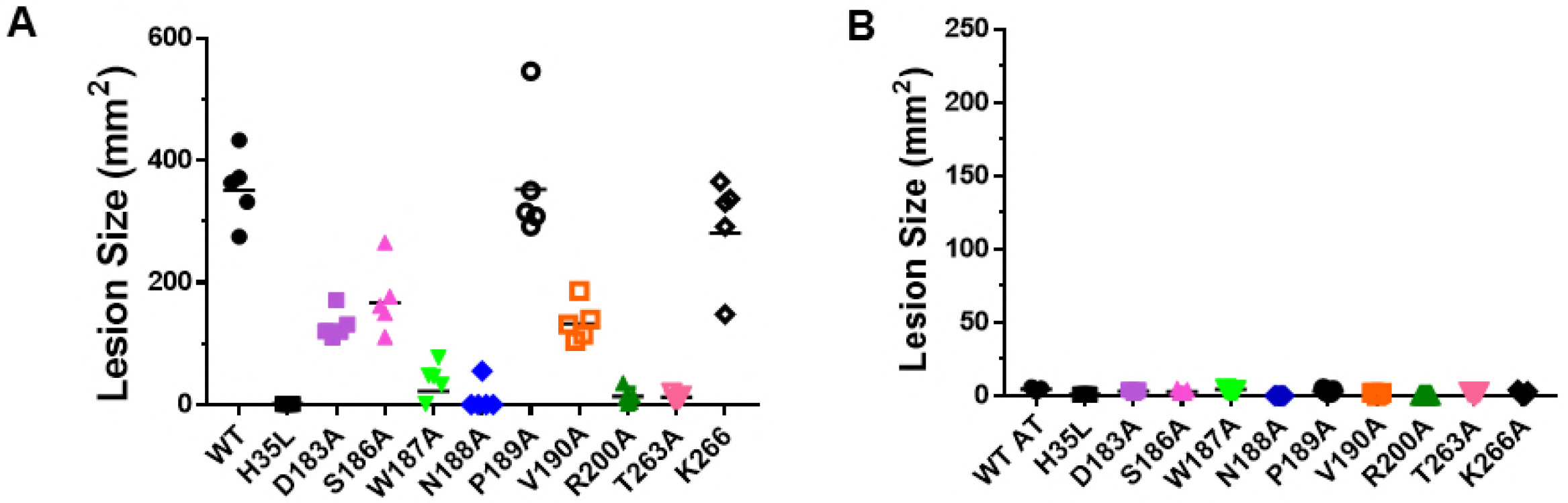
MEDI4893* inhibits purified toxin-induced dermonecrosis. Balb/c mice (n=5) were IP immunized with (A) c-IgG* or (B) MEDI4893* (l0mpk), and ID injected with AT molecules (1 μg). Wild type AT (WT) and AT_H35L_ (H35L)were used respectively as positive and negative controls. Dermonecrotic lesions were measured 24h post ID.

### AT mutations reduce pore formation by decreasing cell binding

AT lyses cells in a multistep process, whereby AT monomers bind ADAM10 on cell membranes, then oligomerize into a heptameric ring and insert into the membrane to form an SDS-resistant transmembrane pore (30). To determine if the lysis defective mutants formed SDS-resistant heptamers in cell membranes, the alanine mutants were incubated with freshly prepared erythrocyte ghost membranes and heptamer formation measured (31) by western blot analysis, as previously described (20). Heptamer formation was measured by densitometry and % heptamer formation was calculated for each mutant relative to WT (**Fig.5**). The oligomerization deficient mutant H35L was included as a negative control. Consistent with the cell lysis assays, N188A, W187A and R200A exhibited the greatest loss in activity, whereas the other mutants lost 50 – 70% of heptamer formation in this assay. These results confirm that the AAs in AT-MEDI4893 contact residues are essential for AT pore formation.

**Fig. 5:**
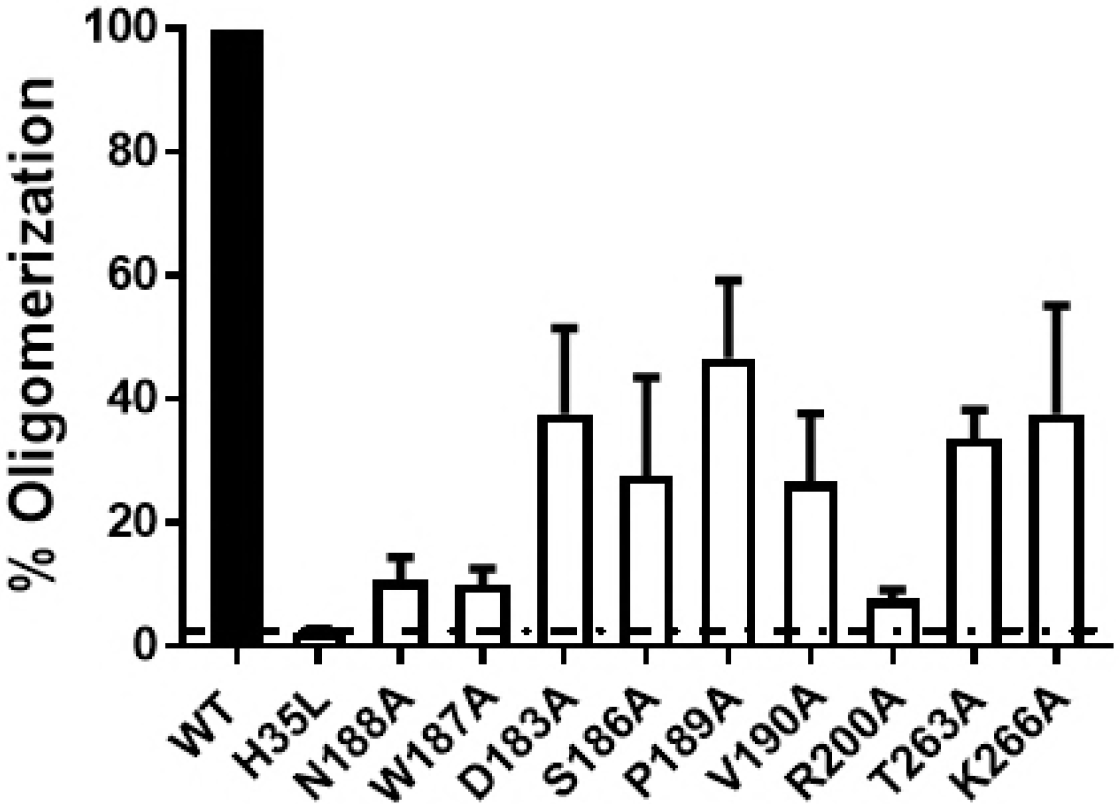
AT mutant heptamer formation. AT mutants were incubated with erythrocyte ghosts at 37°C. Samples were solubilized in SDS-PAGE, and heptamer formation detected by western blot analysis (A). Wild type AT (WT) and AT_H35L_ (H35L)were used respectively as positive and negative controls. Gel is representative of three separate experiments. (B) % of oligomerization was calculated as mean of band intensities from three separate gels. Heptamer formed by WT AT are considered as 100% oligomerization.

Walker et. al. previously reported that amino acid R200 is important for AT binding to cell membranes (31). To determine if the other epitope residues were also important for cell binding, the mutant toxins were biotinylated and binding to A549 cells was measured by flow cytometry (23) and compared with WT-AT and the oligomerization defective mutant, H35L. Similar to the A549 lysis results, AT mutants D183A, P189A, V190A, T263A and K266A exhibited reduced cell binding relative to WT-AT or H35L, whereas no binding was detected with the mutants (W187A, N188A or R200A) most defective for A549 lysis (**Fig. 6, Table S1**), indicating the residues comprising the MEDI4893 epitope are important for cell binding. Consistent with the MEDI4893 neutralization of AT mutant-mediated lysis (**Fig. 3**), the mAb effectively blocked detectable cell binding by all mutants (**Fig. 6**).

**Fig. 6:**
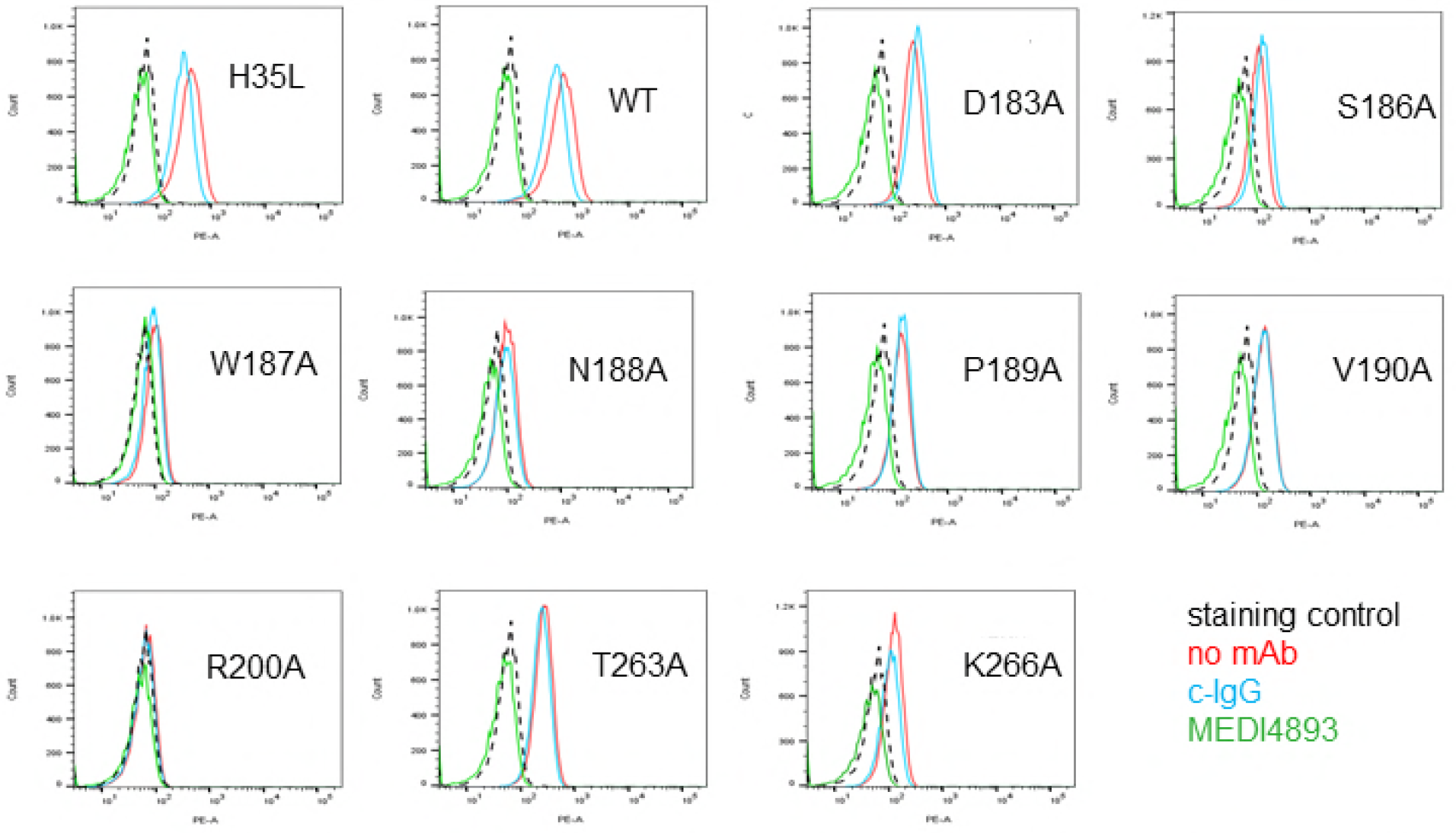
MEDI4893 inhibits AT binding to A549 ceil surface. MEDI4893 (*green*) or negative control IgG (*blue*) were mixed with biotin-conjugated AT WT or mutants, and added to A549 cells. A549 cells were also incubated with toxin alone (*red*). The background consisted of A549 cells alone (*black*). AT binding was measured by cytofluorimetry after addition of streptavidin-FITC

### MEDI4893 binding affinity to AT mutants

The results above indicated MEDI4893 effectively neutralized the lytic alanine mutants *in vitro* and *in vivo*. To further characterize the interaction of MEDI4893 with the epitope mutants, mAb association (k_on_) and dissociation (k_off_) constants for each mutant were measured by surface plasmon resonance and the affinity constant (K_D_) for each variant was calculated. Although the mAb exhibited a modest drop in K_D_ to D183A, S186A, V190A and T263A it retained a sub-nanomolar K_D_, however antibody binding constants to N188A, P189A, R200A and K266A were reduced >10-fold (11- to 26-fold) and no binding to W187A was detected in the assay. These data showed a direct correlation between decreased neutralization of AT mutants in the hemolytic assay and their loss for affinity to MEDI4893 (**Table 1**, **Fig. S2**, r=0.7633) and confirmed that W187 is critical for both MEDI4893 binding and neutralization of AT (**Table 1** **Fig. 3B**).

**Table 1:** Association and dissociation rate constant and apparent binding constants of MEDI4893 to AT alanine mutants. Association (k_on_) and dissociation (k_off_) rate constants were measured using a BIAcore instrument, and apparent binding constant (K_D_) was calculated as k_on_/k_off_

|  | **k<sub>on</sub> (M-1s-1)** | **k<sub>off</sub> (s-1)** | **K<sub>D</sub> (nM)** | **Fold loss** |
| --- | --- | --- | --- | --- |
| **WT** | 1.00E+06 | 1.40E-04 | 0.14 |  |
| **D183A** | 1.78E+06 | 9.08E-04 | 0.511 | 3.5 |
| **S186A** | 7.90E+05 | 1.04E-04 | 0.132 | same |
| **W187A** | No binding observed |  |  |  |
| **N188A** | 2.40E+06 | 5.77E-03 | 2.405 | 16 |
| **P189A** | 2.13E+06 | 3.60E-03 | 1.694 | 11 |
| **V190A** | 2.34E+06 | 6.77E-04 | 0.289 | 2 |
| **R200A** | 1.17E+06 | 3.14E-03 | 2.69 | 18 |
| **T263A** | 1.37E+06 | 2.04E-04 | 0.15 | same |
| **K266A** | 1.35E+06 | 5.15E-03 | 3.812 | 26 |

### MEDI4893* inhibited disease caused by SF8300 strains expressing mutant AT

Although MEDI4893* fully inhibited dermonecrosis induced by ID injection of the nine AT mutants, four mutants (N188A, W187A, R200, K266A) exhibited at least a 15-fold loss of binding by MEDI4893 (**Table 1**). To determine if the mAb prevented disease caused by *S. aureus* strains expressing the mutant toxins with the greatest loss in binding affinity, the alanine mutants were introduced into the USA300 CA-MRSA SF8300 chromosome by allelic exchange. Each mutant was confirmed to express equivalent toxin levels *in vitro* (**Fig. S3**). Balb/c mice (n=10) were passively immunized with MEDI4893* (10mpk) or an isotype control IgG (c-IgG) 24h prior to ID inoculation with each strain (5e7cfu) and lesion sizes were measured 24h post infection. Consistent with the *in vitro* lysis and toxin mediated dermonecrosis results above, the strains expressing N188A, W187A and K266A were less virulent and formed smaller lesions than infection with WT SF8300 and the strain expressing the least lytic mutant R200A was avirulent in this model. Also, MEDI4893* prophylaxis inhibited dermonecrosis caused by the mutant-expressing strains, resulting in lesions resembling infection with an *S. aureus* Δ*hla*. (**Fig. 7** **and Fig. S4**). The AT mutant-expressing strains were similarly avirulent in the murine pneumonia model except for SF8300-N188A which caused disease similar to WT SF8300. Despite a ~16-fold drop in binding affinity for N188A, MEDI4893* prophylaxis significantly reduced death following infection with the N188A mutant strain **(Fig. 8)**. Collectively, these data demonstrate that the AT AAs comprising the MEDI4893-binding epitope are essential not only for AT function, but also for *S. aureus* fitness *in vivo* and that MEDI4893* can neutralize the toxic effects of AT even when its epitope is mutated and its binding affinity reduced.

**Fig. 7:**
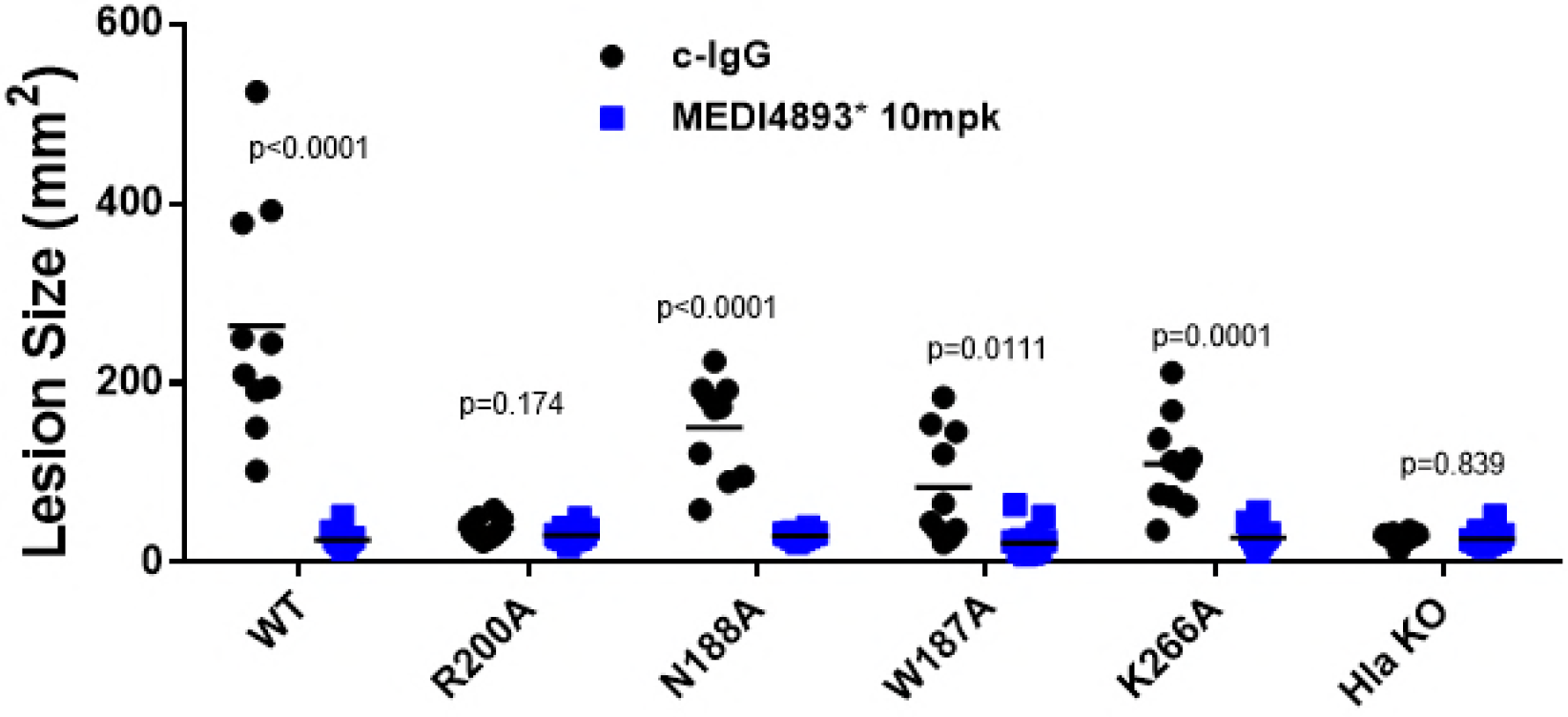
MEDI4893* inhibits USA300 knock-in mutant induced-dermonecrosis. Balb/c mice (n=10) were IP immunized with c-IgG or MEDI4893* (10mpk). and ID injected with SF8300 WT or SF8300 knocking mutants (5e7cfii), Lesion sizes were measured 24hpost infection.

**Fig. 8:**
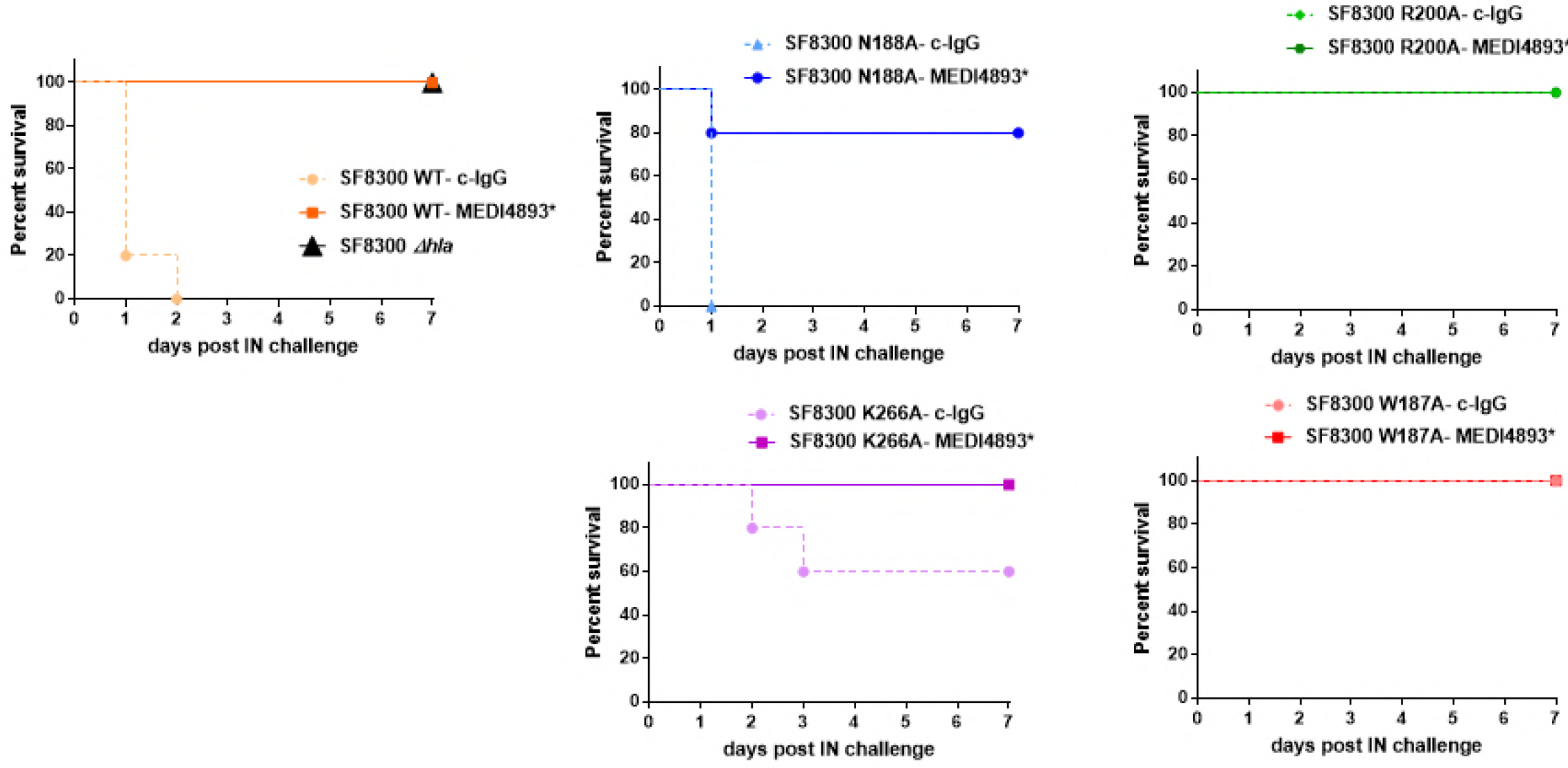
MEDI4893* inhibits USA300 knock-in mutant virulence in pneumonia. C57/B6 mice mice (n=10) were passively immunized with MEDI4893* or c-IgG (15mpk), and IN infected with 1.3 eScfii of SF8300 WT Δ*Hla* or AT mutants. Survival was followed over 6 days post challenge.

## Discussion

The current antibiotic resistance problem has been fueled by widespread use and misuse of broad spectrum antibiotics. Coupled with a greater understanding of the adverse effects that empiric broad-spectrum antibiotic therapy has on the healthy microbiome, there has been an increasing effort to identify alternative strategies to treat infections with antibiotic resistant pathogens. One emerging approach is the use of mAbs to either prevent or treat resistant bacterial infections (1, 32). Although several antibacterial mAb-based treatment strategies are currently in clinical testing, little information is available about resistance to these molecules. Antibacterial childhood vaccines (e.g. diphtheria, tetanus, and pertussis, *Hemophilus influenzae* type b, *Streptococcus pneumoniae*) which rely on a functional antibacterial antibody response have been in use for years or even decades with no apparent emergence of resistance (33). This lack of resistance is likely due in part to the polyclonal antibody response generated by the vaccines and may not predict a similar outcome for an antibacterial mAb.

Unlike antibiotics which kill bacteria and exert direct selective pressure on bacteria, mAb-based antibacterial approaches either neutralize bacterial virulence factors, promote a protective immune response or target the bacteria for opsonophagocytic killing by host immune cells (2, 34). Consequently, the host immune system is responsible for killing the bacteria, not the mAb (35). Since mAbs do not affect bacterial growth directly, it is difficult or even impossible to study the emergence of resistance in vitro or in vivo over the time course of preclinical infection models. To begin to address the questions about circulating resistant *S. aureus* isolates, we conducted two studies in which the AT gene (*hla*) was sequenced from ~1,250 clinical isolates (24–26). In these collections, 58 different AT sequence types were identified and the anti-AT mAb MEDI4893, currently in Phase 2 clinical testing, effectively neutralized all lytic variants. Of these clinical isolates, only 19 encoded an AT protein with a mutation in the MEDI4893 epitope which was effectively neutralized. During Phase 1 study, no AT variants with amino acid substitutions in MEDI4893-binding region were detected despite prolonged exposure of *S. aureus* strains to MEDI4893 due to its extended half-life (26). Because the MEDI4893 epitope is highly conserved and no MEDI4893-resistant mutants were observed thus far, we hypothesized that the AT AAs in the MEDI4893-binding epitope are important for AT function.

In the current study, we mutated 9 AA contact residues in the MEDI4893 epitope to alanine to characterize their roles in AT lytic activity and *S. aureus* fitness and to gain insight of these mutations on MEDI4893 binding and neutralization. Consistent with published results indicating that residue R200 found in the epitope was important for AT cell lysis (31), lytic activity was decreased ≥2-fold in 8 out of 9 mutants constructed herein and of these, 3 mutants lost ≥10-fold of their lytic activity on rabbit RBC or the A549 lung epithelial cell line (Fig.2, Table S1). This loss in toxin activity *in vitro* translated into a similar loss in activity *in vivo* resulting in reduced virulence in both skin and lung infection models. These results support the hypothesis that the MEDI4893 epitope in AT is important not only for AT lytic activity, but also for bacterial fitness in skin and lung infection models. Therefore, it is not surprising that only two MEDI4893 epitope variants (N188T and V190I) were identified in the collection of ~1,250 bloodstream and lung infection clinical isolates (24–26). Although the potential for emergence of MEDI4893 resistance appears to be low, further monitoring of AT sequence variants encoded by circulating clinical isolates is underway that will provide added information regarding AT sequence conservation and the fitness of strains expressing defective or inactive AT variants.

MEDI4893 is an affinity optimized variant of the anti-AT mAb 2A3 (20, 21). When 2A3 was identified, its in *vitro* and *in vivo* neutralizing activity strongly correlated with mAb affinity for the toxin (20). This finding led to an affinity optimization campaign to increase affinity that would hopefully translate into increased potency and protective capacity *in vivo*. Although 2A3 was successfully optimized into MEDI4893 with ~10-fold increased binding affinity (0.60 nM K_D_ for 2A3 and ~0.089 nM K_D_ for MEDI4893) there was no improvement in neutralization activity or protection in preclinical disease models (13, 20, 29). However, the improved MEDI4893 binding affinity may have been beneficial regarding its ability to compensate for epitope mutations in AT. For example, MEDI4893 exhibited 26 and 16-fold losses in affinity for K266A and N188A, respectively, yet it retained a biologically relevant affinity (K_D_ = 3.8 and 2.4 nM, respectively) and neutralizing activity for each mutant was sufficient to prevent disease in the dermonecrosis and pneumonia models. These results indicate that the high affinity binding of MEDI4893 for AT may help overcome a loss in binding due to some mutations in its binding epitope. Taken together with the importance of the MEDI4893 (Suvratoxumab) epitope in AT lytic activity and bacterial fitness, the high mAb affinity may provide an added hurdle for the bacteria on the pathway to develop resistance to AT neutralization by a mAb.

## Materials and methods

### Alpha-toxin alanine mutant expression

The wild-type *hla* gene was PCR-amplified from *S. aureus* SF8300 (USA300) genomic DNA and cloned into a pCN-based *E.coli*-staphylococcal shuttle vector under control of a constitutive promoter based on the *S. aureus clpB* gene promoter (36, 37).

Alanine mutant *hla* expression plasmids were prepared by cloning synthetic DNA fragments containing the mutations into the wild-type expression construct. The alanine mutant plasmids were introduced into *S. aureus* RN4220*Δhla* by electroporation and selected on medium containing 10 μg/mL chloramphenicol. Mutant-expressing strains were cultured overnight at 37°C in BHI (brain heart infusion) broth (Criterion, Inc) with 10 μg/mL chloramphenicol and AT proteins were purified from culture supernatants by cation exchange chromatography using an SP-HP column (GE Healthcare) equilibrated with 30mM Na acetate, pH 5.2, 20mM NaCl, 1mM EDTA and eluted with a linear gradient to 500 mM NaCl.

### Rabbit red blood cell hemolytic assay

Rabbit red blood cell (RBC) hemolytic assay was performed as described (20). Briefly, wild-type AT (WT-AT) or AT mutants were serially diluted two-fold in 50μ1 starting at 10μg/ml, and incubated with 50μ1 of washed RBC (Peel Freeze) for 1h at 37°C. Plates were then centrifuged at 1200rpm for 3min, and 50 μl of supernatant was transferred to new plates. Non-specific human IgG1 R347 was used as negative control (c-IgG) (20). OD_450nm_ was measured with a spectrophotometer (Molecular Devices). Hemolytic activity of each AT mutant was calculated in hemolytic unit per ml (HU/ml) as the inverse dilution corresponding to 50% hemolysis with 10μg/ml corresponding to 1HU/ml.

### A549 cytotoxic assay

Human lung epithelial cell line A549 (ATCC, Manassas, VA) was cultured at 37°C in 5%CO_2_ in Dulbecco’s Modified Eagle Medium (DMEM, VWR International), supplemented with 10% of fetal bovine serum (Gibco) and 2mM glutamine (Invitrogen). Cells were incubated overnight in 96 well plate (VWR International) at 5e4 cells/well. Serial dilutions of AT mutants, or WT-AT was then added to cells for 2h at 37°C, plates were centrifuged at 2000rpm for 2min, and 50μ1 supernatant transferred to new 96 well plate. Cell lysis was measured as release of lactate dehydrogenase (LDH), using the Cytotox 96 nonradioactive assay kit (Promega) following manufacture recommendations. As positive control, cells were lysed with 10% SDS. Background of LDH release was subtracted, and 100% of cell lysis calculated as followed: 100*[(OD_590_ (cells+AT)) / (OD_590_ (Cells +SDS))]

### AT oligomerization on erythrocyte ghosts

Erythrocyte ghosts were prepared as described previously (20) from rabbit blood (Peel Freez). Briefly, 5ml of packed RBC were washed twice in 0.9% NaCl and incubated in 90ml of lysis buffer (5mM Na-phosphate buffer, 1mM EDTA, pH 7.4) overnight at 4°C under constant stirring. Ghosts were obtained after centrifugation at 15,000xg for 20min, washed 3 times with lysis buffer, washed one time in PBS and resuspended in PBS to an optical density at OD_600nm_ 0. 2. Heptamers formed were by incubating 5μl ghosts with purified AT proteins (0.5μg) and PBS in final volume 21 μl for 45min at 37°C. Samples were then solubilized in 7μl of Bolt LDS sample buffer (Invitrogen), incubated for 5min at room temperature and 13 μl was run on 4-12% Bolt MOPS SDS gel (Life technologies). The separated proteins were transferred to nitrocellulose membrane in Bolt-transfer buffer (Invitrogen) with 10% methanol for overnight at 15V, blocked with Odyssey blocking buffer (LI-COR) for 1h, probed with rabbit anti-AT IgG (2μg/ml) for 2h at room temperature. The AT bands were detected after 1h incubation with IRDye 680RD-conjugated-donkey anti-rabbit IgG (LI-COR) by Odyssey--fluorescence imager (LI-COR). Band intensities were calculated with Odyssey Image Studio Lite software.

### MEDI4893 affinity to AT mutants

Kinetic rate constants (*k*_on_ and *k*_off_) for binding of the MEDI4893 to WT-AT, and each mutant were measured by employing an IgG capture assay on a BIAcore T200 instrument. Protein A was immobilized on a CM5 sensor chip with a final surface density of ~ 2,000 resonance units (RUs). MEDI4893 was prepared at 10nM in instrument buffer (HBS-EP buffer; 0.01MHEPES, pH 7.4, 0.15MNaCl, 3mMEDTA, and 0.005% P-20), along with 2-fold serial dilutions of AT (0.048 nM to 50 nM). A sequential approach was utilized for kinetic measurements. MEDI4893 was first injected over the capture surface, at a flow rate of 10μl/min. Once the binding of the captured IgG stabilized, a single concentration of the AT protein was injected over both capture and reference surfaces, at a flow rate of 50 μl/min. The resulting binding response curves yielded the association phase data. Following the injection of AT, the flow was then switched back to instrument buffer for 10min to permit the collection of dissociation phase data, followed by a 1min pulse of 10mM glycine, pH 1.7 to regenerate the Protein A surfaces on the chip. Binding responses from duplicate injections of each concentration of AT were recorded against anti-AT mAb MEDI4893. In addition, several buffer injections were interspersed throughout the injection series. Select buffer injections were used along with the reference cell responses to correct the raw data sets for injection artifacts and/or nonspecific binding interactions, commonly referred to as “double referencing”. Fully corrected binding data were then globally fit to a 1:1 binding model (BIAevaluation 4.1 software; BIAcore, Inc.) that included a term to correct for possible mass transport-limited binding. These analyses determined the kinetic rate constants *k*_on_ and *k*_off_, from which the apparent dissociation constant (*K*D) was calculated as *k*_off_/*k*_on_.

### Alpha toxin binding by cytofluorimetry

AT binding to A549 cells, and inhibitory effect of MEDI4893 was measured as previously described (23) with some modifications. Briefly, AT mutants or WT were biotinylated using the EZ link Sulfo-NHS-LC biotinyaltion kit (Thermo Fisher Scientific) following manufacturer protocol. All incubations and washes were performed in FACS buffer (PBS, 0.5% BSA, 0.1% Tween). AT and MEDI4893* were pre-incubated for 30min at room temperature at a molar ratio 1:5. Cells were first blocked in with human Fc blocker (eBioscience), then incubated for 1h at 4°C with AT alone or MEDI4893-AT mix. Following one wash, cells were then incubated for 30min at 4°C with Streptavidin FITC conjugate (eBioscience). After two washes, AT binding was then quantified with a LSRII flow cytometer (BD Biosciences) and data analyzed with Flow Jo Software (Tree Star, Inc., Ashland, OR).

### Chromosomal allelic exchange of *hla* alanine mutants in strain SF8300

Alanine mutation-containing sequences were subcloned into temperature sensitive allelic exchange vector pBD100 (Binh Diep, UCSF). Constructs were transferred from strain RN4220 into strain SF8300 by phi11 transduction. Chromosomal plasmid integrants were obtained by temperature selection and verified by PCR. Subsequent negative selection for plasmid excision was performed by plating on media containing anhydrotetracycline. Alanine substitutions in the chromosomal *hla* gene were verified by PCR and sequencing.

### Mouse dermonecrosis

Demonecrosis studies were conducted as previously described (20). Six-week-old female Balb/c mice (Harlan) were passively immunized with MEDI4893* (10mpk) or c-IgG (20), and intradermally (ID) challenged 24h later with SF8300 WT, or SF8300 expressing AT alanine mutants (5e7cfu). Dermonecrosis was also induced by ID injection of purified alanine mutants or WT (diluted in cold PBS at 1μg/50μl). Lesion sizes were measured 24h and 7 days after infection.

### Mouse pneumonia

Lethal pneumonia was induced as reported previously (13). Six-week-old female C57/B6 mice (Jackson) were passively immunized with MEDI48938 (15mpk) or c-IgG, and intra-nasally infected 24h later with SF8300 WT or SF8300 expressing AT alanine mutants (1.5e8cfu in 50μl). Animal survival was monitored over 6 days post infection.

### Mouse models

All experiments were performed in accordance with institutional guidelines following experimental protocol review and approval by the Institutional Biosafety Committee (IBC) and the Institutional Animal Care and Use Committee (IACUC) at MedImmune.

## Figure legends

**Fig. 1: Interface between MEDI4893 Fab HC (green) and AT (pink) (A) and MEDI4893 Fab LC (purple) and AT (pink) (B).** Both chains of the Fab interact with AT and create hydrogen bonds (dotted lines). Residue W187 interacts with the heavy chain (HC) through hydrogen bound, and with W32 on light chain (LC) by π–π stacking interaction.

**Fig. 2: AT mutant lytic activity on rabbit RBC (A) and human A549 cell line (B). (A)** Washed RBCs were incubated with serial dilutions of WT-AT or AT mutants (10 to 0.005μg/ml). Hemolysis was measured by the amount of hemoglobin released in the supernatant, and calculated as follows: 100*[(OD_AT_)/(OD_SDS_)]. (B) A549 cells were incubated with serial dilutions of WT-AT or AT mutants (20 to 0.01μg/ml). LDH was measured in supernatants after two-hour incubation at 37°C. *%* cytolysis was calculated as follows: 100*[100-(OD_AT_) / (OD_SDS_)].

**Fig. 3: MEDI4893 neutralization in vitro on rabbit RBC hemolysis (A) and A549 lysis (B)** WT or mutant AT was mixed with MEDI4893 serial dilutions with RBC (0.1μg/ml) (A) or with (B) A549 cells (20μg/ml). % hemolysis or cell lysis inhibition was calculated as follows: 100*[100-(OD_AT+mAb_) / (OD_AT alone_)]. Data are representative of three independent experiments for AT: MEDI4893 ratio 1:2.

**Fig. 4: MEDI4893* inhibits purified toxin-induced dermonecrosis.** Balb/c mice (n=5) were immunized intraperitoneally with 10mpk (A) c-IgG or (B) MEDI4893*, and injected intradermally with WT or mutant AT (1μg). AT_H35L_ (H35L) was included as a negative control. Dermonecrotic lesions were measured 24h post-infection.

**Fig. 5: AT mutant heptamer formation.** The alanine mutants were incubated with erythrocyte ghosts at 37°C. Samples were solubilized in SDS-PAGE, and heptamer formation detected by western blot analysis (A). WT and H35L-AT were used as positive and negative controls. The blot is representative of three independent experiments. (B) % oligomerization was calculated from mean band intensities on three separate blots. Heptamers formed by WT-AT are considered as 100% oligomerization.

**Fig. 6: MEDI4893 inhibits AT binding to A549 cell surface.** Biotin-conjugated WT or mutant AT was incubated with A549 cells in the presence of MEDI4893 (green) or c-IgG (blue). A549 cells were also incubated with toxin alone (red). The background consisted of A549 cells alone (black). AT binding was measured by cytofluorimetry after addition of streptavidin-FITC

**Fig. 7: MEDI4893* inhibits USA300 knock-in mutant induced-dermonecrosis.** Balb/c mice (n=10) were immunized intraperitoneally with c-IgG or MEDI4893* (10mpk) 24 h prior to intradermal infection with *S. aureus* SF8300 or the SF8300 *hla* knock-in mutants (5e7cfu). Lesion sizes were measured 24h post infection.

**Fig. 8: MEDI4893* inhibits USA300 knock-in mutant virulence in pneumonia.** C57/B6 mice mice (n=10) were passively immunized with MEDI4893* or c-IgG (15mpk), and infected intranasally with 1.3e8cfu of WT, Δ*hla* or *hla* knock-in mutant SF8300. Survival was followed over 6 days post-challenge.

